# A double-blinded, placebo-controlled field trial of an OspA-based oral reservoir targeted vaccine against *Borrelia burgdorferi*

**DOI:** 10.64898/2026.04.21.719976

**Authors:** Amy M. Schwartz, Ferney Henao-Ceballos, Kathryn Arnold, Julia Poje, Max Waugh, Greg Joyner, Jose F. Azevedo, Tyler Baccam, Eric Kontowicz, Kurayi Mahachi, Paige Witucki, Suman Kundu, Nisha Nair, Felix Pabon-Rodriguez, Grant Brown, Maryland Field Team, Christine Petersen, Maria Gomes-Solecki

## Abstract

Lyme disease, caused by *Borrelia burgdorferi* and transmitted by *Ixodes scapularis* ticks, remains a significant vector-borne illness in the United States. Small mammal reservoirs, particularly *Peromyscus leucopus*, play a critical role in *B. burgdorferi* maintenance. Here we conducted a five-year, randomized, double-blinded, placebo-controlled field trial deploying an oral OspA-based reservoir targeted vaccine (RTV) across seven Maryland sites. Bayesian modeling provided estimates of vaccine impact on mouse anti-OspA antibody levels, nymphal tick infection prevalence (NIP), mouse infection rates, and seroconversion to *B. burgdorferi* in hunting dogs. RTV sites exhibited an estimated 10.5% proportional increase in protective murine anti-OspA antibody levels and a 15.4% reduction in NIP by year five. We also found a lower infection prevalence in mouse blood fed nymphal ticks (9.8%). RTV sites exhibited modest decreases in mouse infection prevalence and dog seroconversion rates were similar between groups. Our results indicate that anti-OspA antibody in vaccinated-infected *P. leucopus* reduced *B. burgdorferi* summertime larval infection prevalence, measured as NIP reductions the following spring. This suggests that OspA-based oral RTV reduces *B. burgdorferi* transstadial transmission within tick populations. Our findings advance development of reservoir targeted solutions for Lyme disease prevention. Further evaluation of impacts on incidental hosts is needed.

## Introduction

Lyme disease is the most reported vector-borne disease in the United States. An estimated 475,000 Lyme disease diagnoses occur annually^1^. While Lyme disease risk has historically been concentrated in the Northeast, Mid-Atlantic, and Upper Midwest, in recent years, risk has expanded to adjacent regions with suitable habitat^2,3^.

Lyme disease is caused by the bacteria *Borrelia burgdorferi* and can be transmitted to humans through the bite of an infected *Ixodes* spp tick^4^. *Ixodes scapularis* ticks primarily acquire *B. burgdorferi* from small mammal hosts while feeding during the larval and nymphal stages^5^. *Peromyscus leucopus* (the white-footed mouse) is the primary reservoir host of *B. burgdorferi* in the Northeastern United States^6^. Nymphal ticks are the largest contributors to human Lyme disease due to their abundance during months of peak human outdoor activity and their small size, which hinders detection and removal. Both nymphal infection prevalence and density of infected nymphs are important indicators of Lyme disease risk^7–9^.

Studies have shown that most tick exposures occur in privately owned, peridomestic settings^10,11^. As such, interventions that reduce the prevalence of ticks infected with *B. burgdorferi* on peridomestic and recreational property hold high potential. A reservoir targeted vaccine (RTV) intervention could potentially disrupt multiple components of the *B. burgdorferi* enzootic cycle^12^ through oral vaccination of *P. leucopus*^13^ and other rodent reservoir hosts. The RTV tested in this field study is based on outer surface protein A (OspA) of *B. burgdorferi* expressed on inactivated cells of recombinant *E. coli*^9^. A similar formulation received USDA license approval in 2023. The study was conducted in Maryland at seven privately owned field sites (Figure 1) over a 5-year period (2020-2024). Previous studies evaluating both live^13^ and inactivated^14^ RTV over multiple years demonstrated significant impacts on infection prevalence of larvae and nymphal *I. scapularis* collected from mice and by drag sampling^13,14^. Our hypothesis for this study was that oral bait vaccination of *P. leucopus* reservoir hosts with an OspA based RTV in forested regions of MD would significantly reduce *B. burgdorferi* burden in nymphal ticks. In addition, we evaluated the RTV effect on other components of the enzootic cycle including mouse infection prevalence and engorged nymphal tick infection prevalence; importantly, we assessed hunting dogs as proxy indicators for human Lyme disease risk. Hunting dogs have been shown to have high exposure to ticks and tick-borne pathogens^15^. Thus, we leveraged the presence of hunting kennels situated on field sites to evaluate the impact of RTV on hunting dog *B. burgdorferi* seroconversion^16^. The primary outcome of this study was to evaluate if deployment of the inactivated version of the RTV would reproduce the reductions in nymphal infection prevalence (NIP) observed in a previous field trial^13^, in distinct geographic location (MD), to understand RTV impact on Lyme disease risk outside New England and southern New York. We repeated critical parts of the original study design to compare measurement of OspA seroprevalence in mice after RTV deployment and determine *B. burgdorferi* infection prevalence in questing nymphs, and we added the following modifications: the three assigned RTV sites were treated with vaccine during 5 years (in the first study, only 1 of 4 RTV assigned sites received vaccine for 5 years^13^); and two-thirds (2/3) of the RTV was distributed through automatic carousels and one-third (1/3) was distributed in Sherman traps.

**Figure 1.**
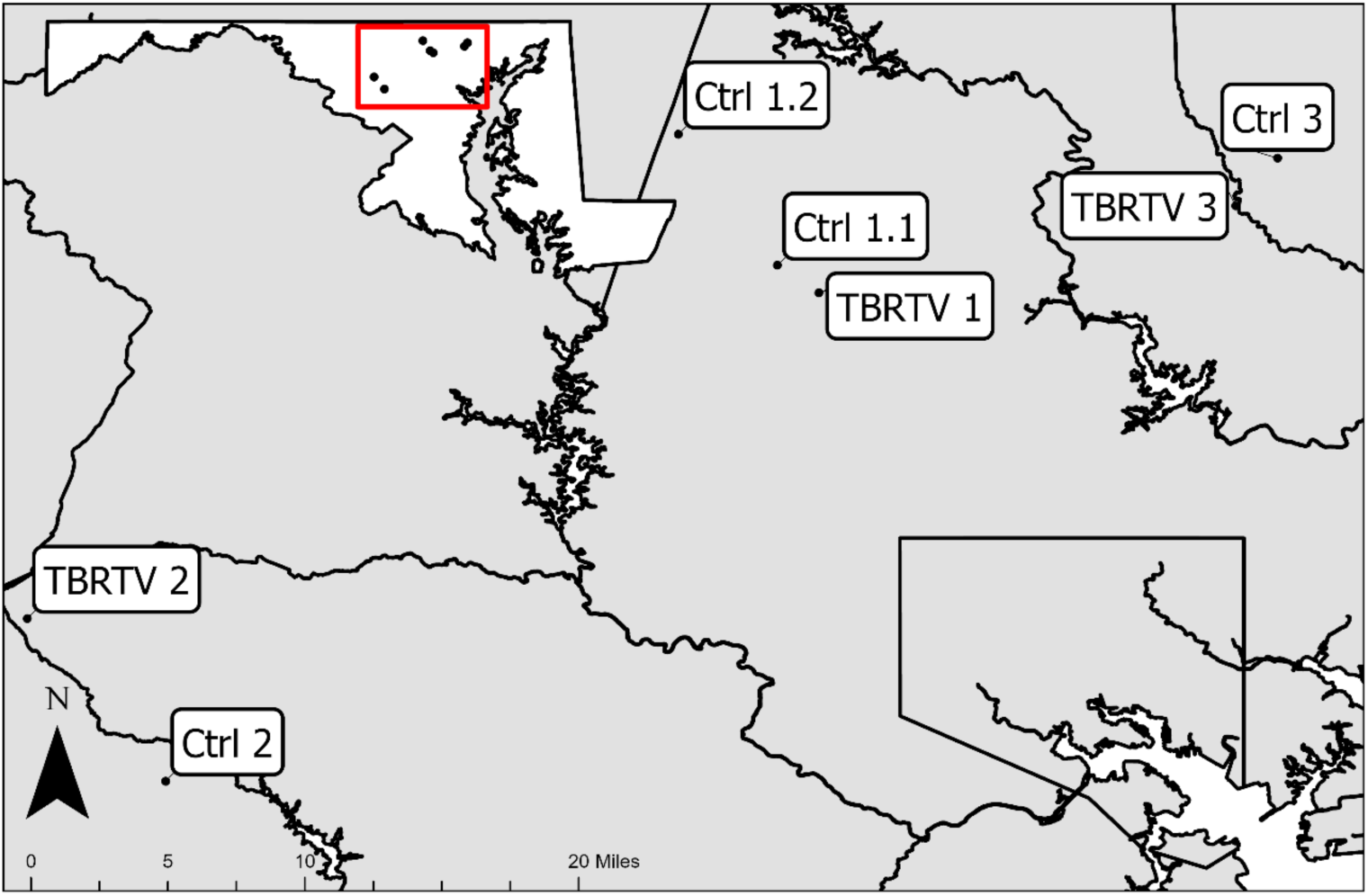
Locations of RTV and Ctrl field sites in the Mid-Atlantic state of Maryland. Sites were located in Harford, Baltimore, Montgomery, and Howard Counties. Ctrl 1.1 was active during years 1 and 2 and Ctrl 1.2 was active during years 3-5.

## Results

### Quality control of laboratory produced inactivated RTV bait

Each lot of RTV bait was assessed for immunogenicity and vaccine efficacy. Spectrophotometric analysis of BPL-treated (OD_600_=0) versus BPL-untreated (OD_600_>1.0) *E. coli* expressing *B. burgdorferi* OspA confirmed inactivation of the RTV bait, and OspA antigen expression was confirmed by immunoblot using anti-OspA monoclonal antibody LA2.2. Determination of serologic presence of anti-OspA IgG in mice was done by ELISA. RTV bait-vaccinated mice developed anti-OspA antibodies above the predicted protective threshold cutoff, in contrast to the controls. Analysis of *B. burgdorferi* dissemination three weeks after tick challenge showed that none of the mice that received RTV bait developed antibodies to *B. burgdorferi* pepVF and did not have *B. burgdorferi flaB* DNA in heart tissue, in contrast to the control.

### Nymphal Infection Prevalence Outcomes

A total of 831 flat, nymphal *Ixodes* ticks were tested over the study period. In the first year of the study (baseline, Year 1, 2020), the unadjusted observed average NIP among RTV sites was higher (22.2%) than that of Ctrl sites (6.5%); however, the difference was not statistically significant (p-value: 0.052). Average annual NIP ranged from 10.2 to 22.2% among RTV sites and from 4.6 to 16.2% among control sites (Table 1). With respect to multiplicative percent change in NIP from baseline to year 5, RTV sites had a 42.6% decrease in NIP (22.2% to 12.7%) and Ctrl sites had a 148.3% increase (6.5% to 16.2%) with fluctuations among RTV and Ctrl groups over the study period. Testing the additive raw difference at year five versus year one between the RTV and Ctrl groups using a normal proportion test approach gives an estimated 19.2% difference in the relative decrease in NIP with a p=0.038 (Figure 2A).

**Table 1.**
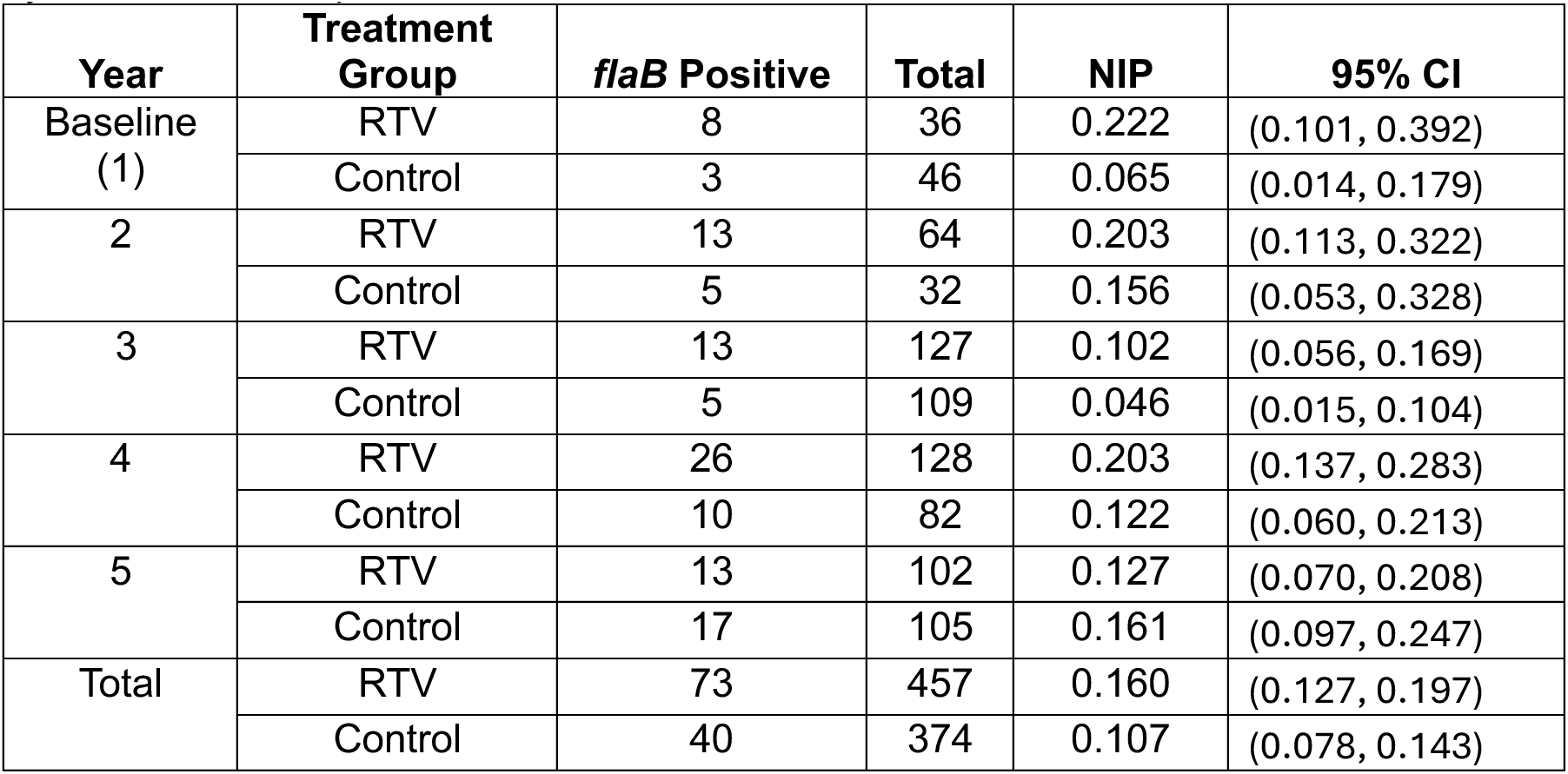
Unadjusted observed flat Nymphal Ixodes (collected by drag) Infection Prevalence by Treatment Group and Year.

| Year | Treatment Group | *flaB* Positive | Total | NIP | 95% CI |
| --- | --- | --- | --- | --- | --- |
| Baseline (1) | RTV | 8 | 36 | 0.222 | (0.101, 0.392) |
|  | Control | 3 | 46 | 0.065 | (0.014, 0.179) |
| 2 | RTV | 13 | 64 | 0.203 | (0.113, 0.322) |
|  | Control | 5 | 32 | 0.156 | (0.053, 0.328) |
| 3 | RTV | 13 | 127 | 0.102 | (0.056, 0.169) |
|  | Control | 5 | 109 | 0.046 | (0.015, 0.104) |
| 4 | RTV | 26 | 128 | 0.203 | (0.137, 0.283) |
|  | Control | 10 | 82 | 0.122 | (0.060, 0.213) |
| 5 | RTV | 13 | 102 | 0.127 | (0.070, 0.208) |
|  | Control | 17 | 105 | 0.161 | (0.097, 0.247) |
| Total | RTV | 73 | 457 | 0.160 | (0.127, 0.197) |
|  | Control | 40 | 374 | 0.107 | (0.078, 0.143) |

**Figure 2.**
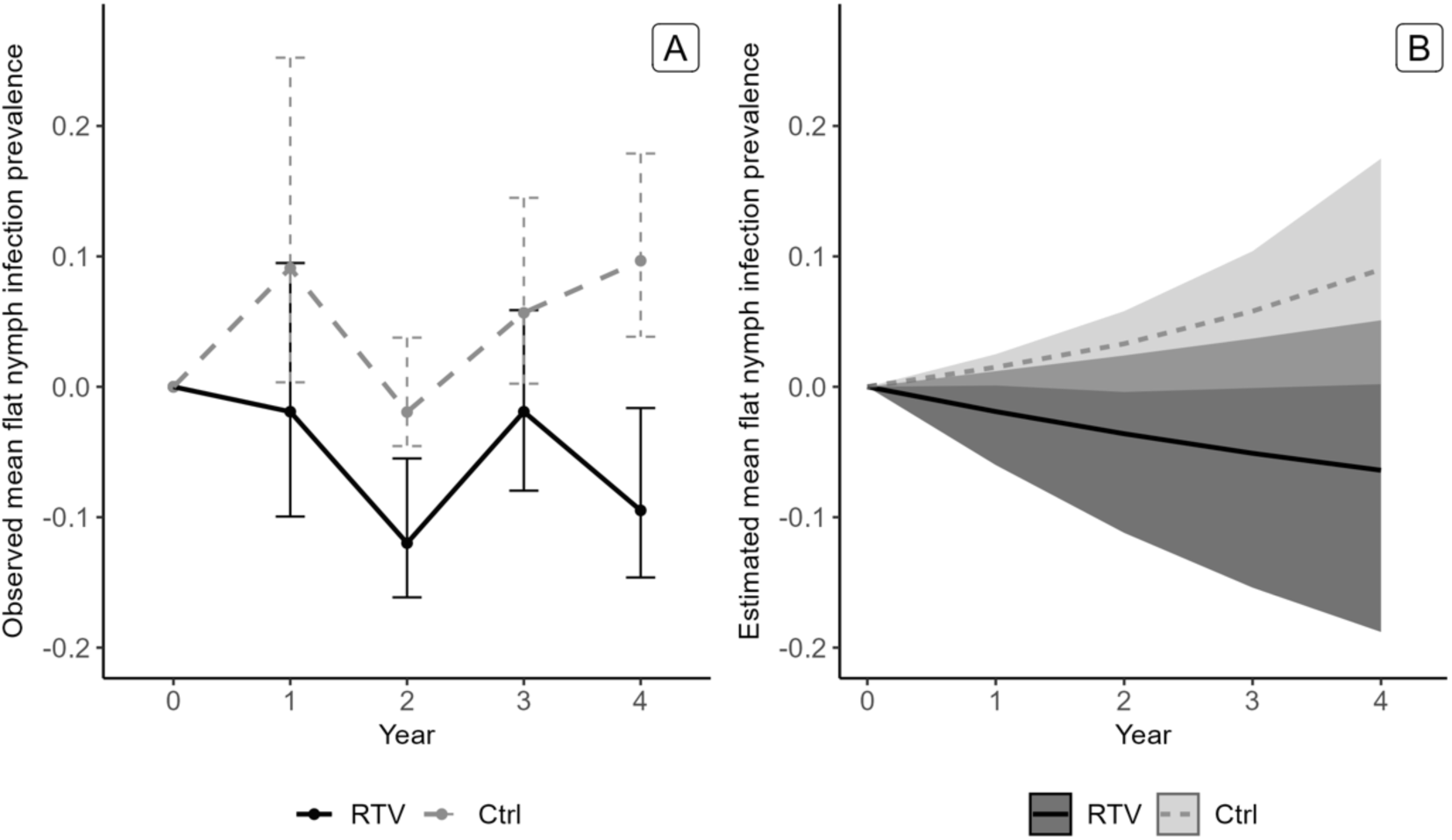
Flat nymphal tick infection prevalence with B. burgdorferi by year from baseline. A. Unadjusted observed proportion including 95% confidence intervals; B. Estimated mean generated from GLMM posterior samples; the shadowing represents 95% Bayesian credible intervals.

All parameters of the GLMM achieved convergence with a value of ∼1 for the Gelman-Rubin statistic. The mean NIP estimates generated from the posterior samples of the model suggested that NIP decreased among RTV sites from baseline (Figure 2B; Table 2). Comparing each year with baseline, the estimated mean effect of the RTV increased consistently from 0.034 to 0.154 and the probability that the RTV resulted in NIP reductions ranged from 0.974 to 0.981. The results of the model suggest a 15.4% (Cr-I: 0.9%, 30.5%) reduction in NIP from baseline among RTV sites in the final year of the study compared with Ctrl sites.

**Table 2.**
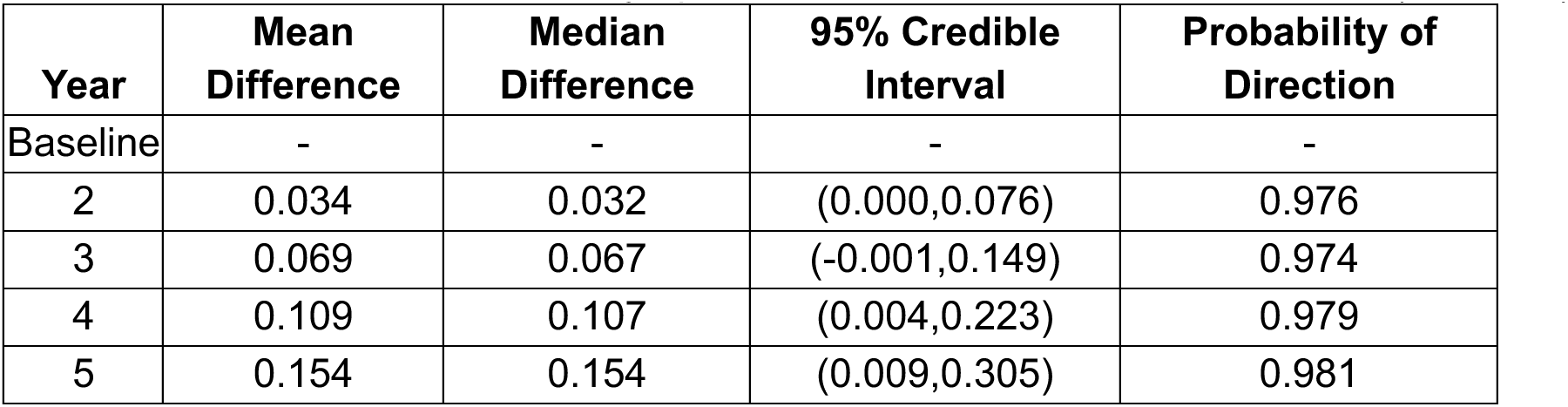
Estimated Effect on Flat Nymphal Ixodes Tick Infection from Baseline (Ctrl-RTV)

| Year | Mean Difference | Median Difference | 95% Credible Interval | Probability of Direction |
| --- | --- | --- | --- | --- |
| Baseline | - | - | - | - |
| 2 | 0.034 | 0.032 | (0.000,0.076) | 0.976 |
| 3 | 0.069 | 0.067 | (-0.001,0.149) | 0.974 |
| 4 | 0.109 | 0.107 | (0.004,0.223) | 0.979 |
| 5 | 0.154 | 0.154 | (0.009,0.305) | 0.981 |

### Mouse Outcomes

Overall, 7,981 mouse capture events occurred with a total of 1,618 unique *Peromyscus* spp. mice captured and tagged over the course of the study. A total of 242 mouse capture events were missing a tag with an identification number. Mice were recaptured a median of three times with a range of 1 to 34 recapture events. Of mice with recorded sex, approximately 58% of captured mice were male and 42% were female. Mice from RTV sites did not significantly differ from Ctrl control sites by age group or sex. Less than 10% of mice were recaptured the next season, thus ∼ 90% of *Peromyscus* in the study lived one year.

### Mouse OspA Seroprevalence

A total of 2,483 mouse blood samples were tested for anti-OspA antibodies. Except for the baseline year (2020, year 1), we measured significantly increased seroconversion to OspA in serum from *P. leucopus* trapped in RTV sites as compared to Ctrl sites (Figure 3). Across years, mean OD_450_ was 0.51 among mice captured from RTV sites compared with 0.39 among mice captured from Ctrl sites (p<0.0001). 15.3% (Range: 6.0-24.2%) of samples tested above the predicted correlate of protection threshold (>0.800) compared with 5.4% (Range: 2.7-9.5%) from control (Ctrl) sites (p-value<0.01) (Figure 4A, Table 3).

**Figure 3.**
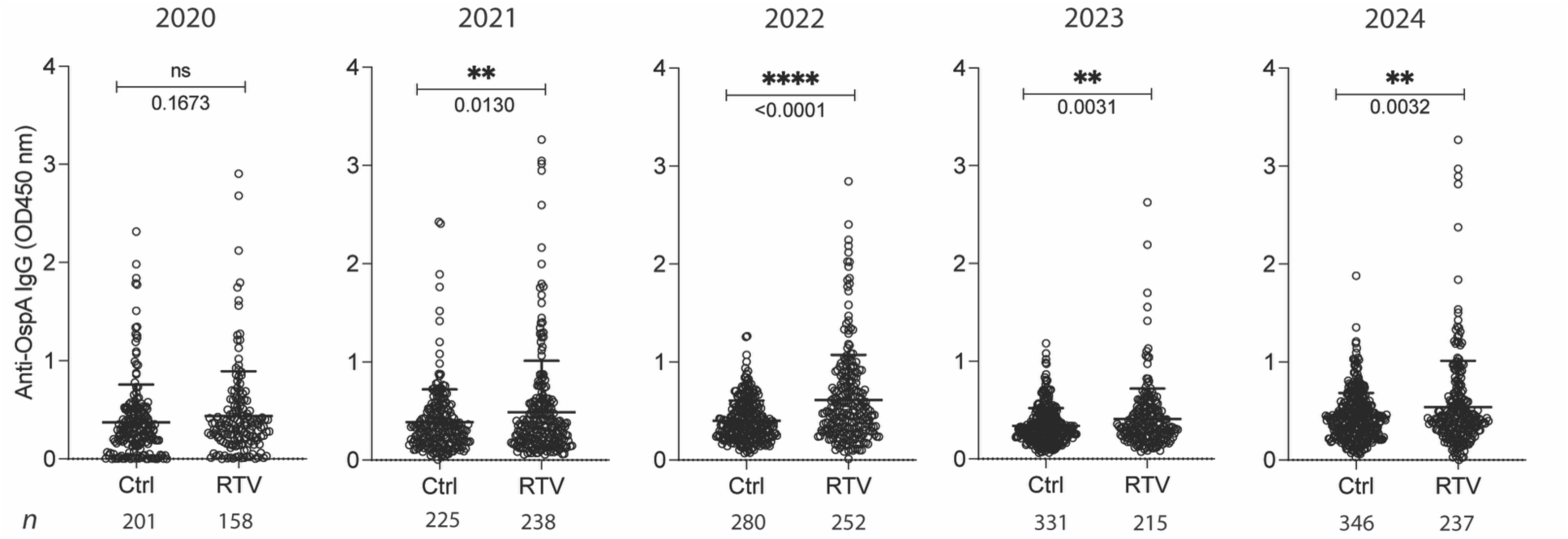
Antibody (IgG) seroconversion to OspA in serum from Peromyscus leucopus captured in the MD field sites between May and July of 2020 to 2024. Scatter dot plots representing antibody to B. burgdorferi OspA measured by ELISA OD_450_ (mean with SD). Legend: n, number of samples tested, Ctrl, received untreated pellets; RTV, received pellets coated with inactivated E. coli expressing OspA. Statistics by Unpaired t test with Welch’s correction.

**Figure 4.**
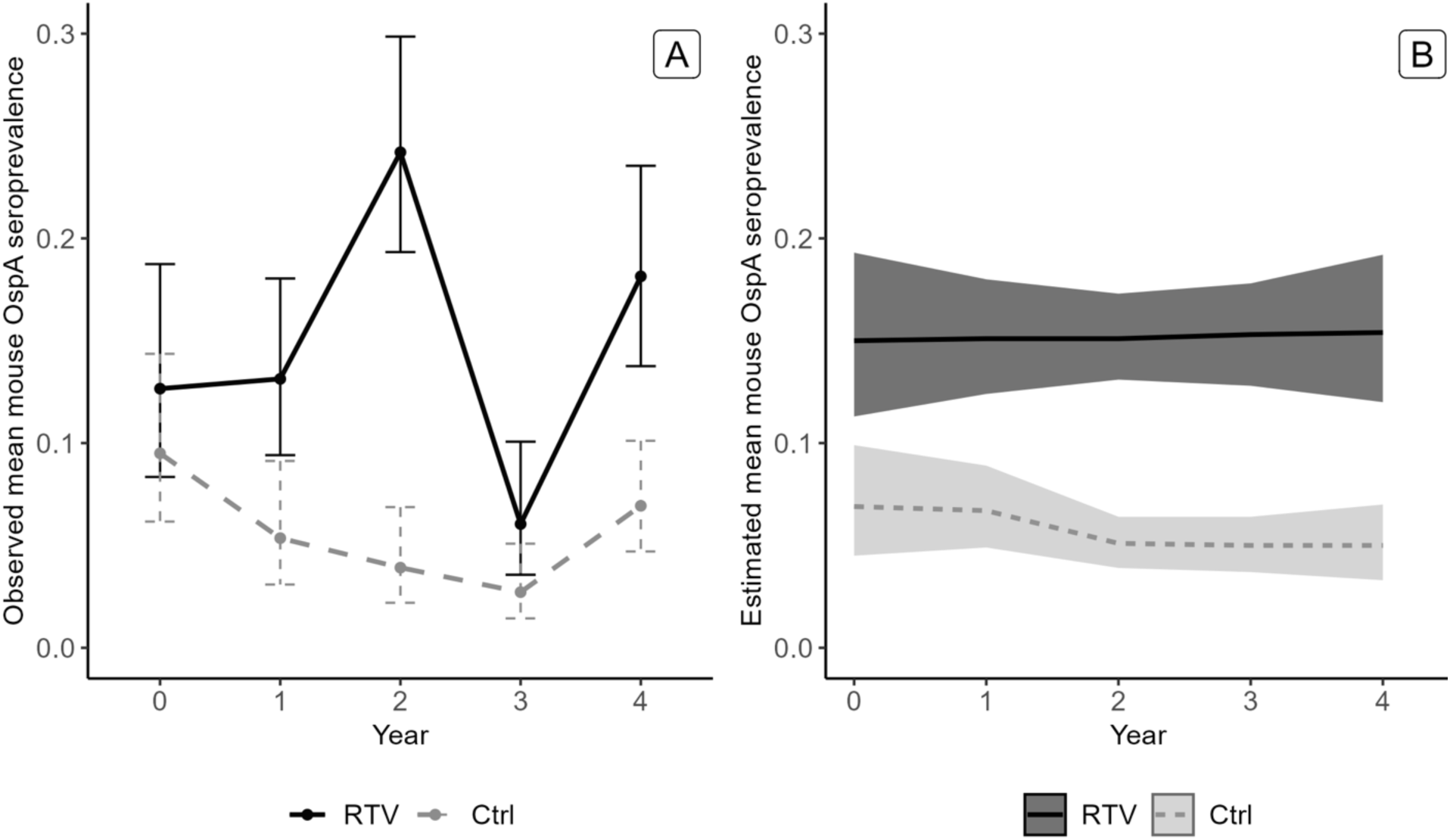
Mouse protective OspA seroprevalence by year. A. Unadjusted observed proportion including 95% confidence intervals; B. Estimated mean generated from GLMM posterior samples; the shadowing represents 95% Bayesian credible intervals.

**Table 3:**
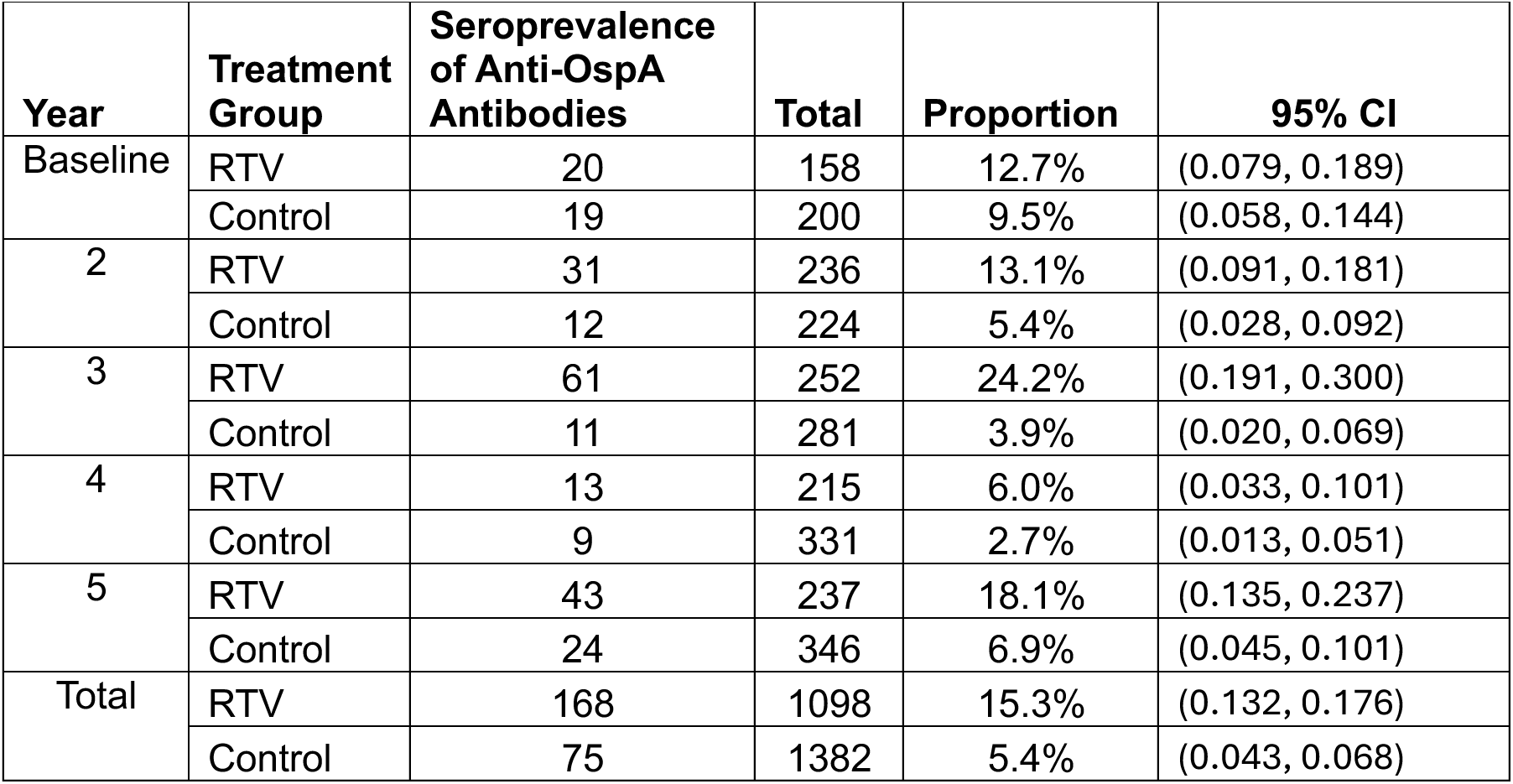
Unadjusted observed mouse OspA Seroprevalence above the predicted protective threshold by Treatment Group and Year.

| Year | Treatment Group | Seroprevalence of Anti-OspA Antibodies | Total | Proportion | 95% CI |
| --- | --- | --- | --- | --- | --- |
| Baseline | RTV | 20 | 158 | 12.7% | (0.079, 0.189) |
|  | Control | 19 | 200 | 9.5% | (0.058, 0.144) |
| 2 | RTV | 31 | 236 | 13.1% | (0.091, 0.181) |
|  | Control | 12 | 224 | 5.4% | (0.028, 0.092) |
| 3 | RTV | 61 | 252 | 24.2% | (0.191, 0.300) |
|  | Control | 11 | 281 | 3.9% | (0.020, 0.069) |
| 4 | RTV | 13 | 215 | 6.0% | (0.033, 0.101) |
|  | Control | 9 | 331 | 2.7% | (0.013, 0.051) |
| 5 | RTV | 43 | 237 | 18.1% | (0.135, 0.237) |
|  | Control | 24 | 346 | 6.9% | (0.045, 0.101) |
| Total | RTV | 168 | 1098 | 15.3% | (0.132, 0.176) |
|  | Control | 75 | 1382 | 5.4% | (0.043, 0.068) |

Results from the GLMM show that an estimated larger proportion of mice from RTV sites had protective levels of anti-OspA antibodies compared with Ctrl sites. Starting in the baseline year, the mean difference between Ctrl and RTV sites was 0.081 (Cr-I: 0.034,0.130); the mean estimated RTV effect increased consistently each year reaching a mean difference of 0.105 (Cr-I: 0.065,0.146) (Figure 4B; Table 4) in the final year. The probability that the RTV resulted in a positive effect was approximately 1.0 across all 5 years, which is very strong evidence that anti-OspA antibodies in RTV sites were associated with a protective effect.

**Table 4.** Estimated Effect on mouse OspA Seroprevalence above the predicted protective threshold from Baseline (Ctrl-RTV)

| Year | Mean Difference | Median Difference | 95% Credible Interval | Probability of Direction |
| --- | --- | --- | --- | --- |
| Baseline | 0.081 | 0.081 | (0.034,0.130) | 1.000 |
| 2 | 0.083 | 0.083 | (0.049,0.118) | 1.000 |
| 3 | 0.101 | 0.101 | (0.076,0.126) | 1.000 |
| 4 | 0.103 | 0.103 | (0.075,0.131) | 1.000 |
| 5 | 0.105 | 0.104 | (0.065,0.146) | 1.000 |

### Mouse exposure to B. burgdorferi and infection

A total of 2,440 mouse blood specimens were tested by pepVF ELISA to assess exposure to *B. burgdorferi*. The unadjusted observed proportions of positive specimens among RTV and Ctrl sites fluctuated among years and ranged from 32.9% (baseline) to 68.7% (year three) among RTV sites, and 39.0% (baseline) to 61.4% (year three) among Ctrl sites; overall proportions did not differ between RTV and Ctrl sites (Supplementary Table 1).

A total of 2,760 ear tissue samples were tested by qPCR to determine presence of *B. burgdorferi* spirochetes. Overall, spirochetes were detected in approximately 40% of mouse ear tissue samples with proportions ranging from 27.8% to 49.5% among years; unadjusted observed proportions did not differ by RTV (Overall: 39.7%; Range: 28.2-47.8%) and Ctrl groups (Overall: 40.7%; Range: 25.2-51.4%) (Table 5). However, the proportion of mice with detectable spirochetes in ear tissue increased from baseline at both RTV (Percent change: 66.4%) and Ctrl sites (Percent change: 67.3%) (Figure 5A),

**Figure 5.**
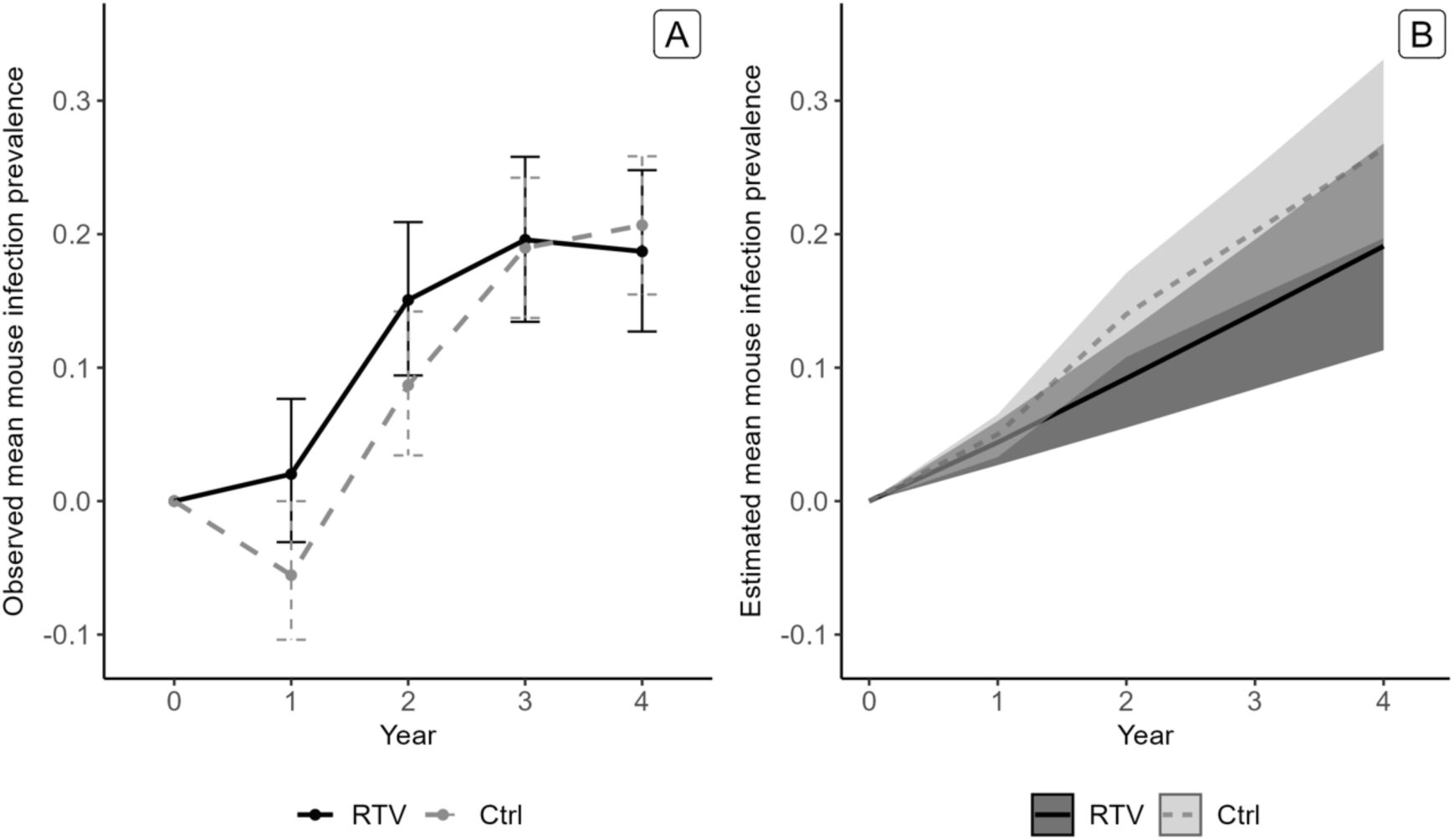
Mouse ear infection prevalence with B. burgdorferi (qPCR) by year from baseline. A. Unadjusted observed proportion including 95% confidence intervals; B. Estimated mean generated from GLMM posterior samples; the shadowing represents 95% Bayesian credible intervals.

**Table 5.** Unadjusted observed detection of B. burgdorferi (qPCR) in Mouse Ear Biopsies by Treatment Group and Year.

| Year | Treatment Group | *flaB* Positive | Total | Proportion | 95% CI |
| --- | --- | --- | --- | --- | --- |
| Baseline | RTV | 53 | 188 | 28.2% | (0.219, 0.352) |
|  | Control | 71 | 231 | 30.7% | (0.249, 0.371) |
| 2 | RTV | 84 | 278 | 30.2% | (0.249, 0.360) |
|  | Control | 67 | 266 | 25.2% | (0.201, 0.309) |
| 3 | RTV | 122 | 282 | 43.3% | (0.374, 0.493) |
|  | Control | 123 | 312 | 39.4% | (0.340, 0.451) |
| 4 | RTV | 118 | 247 | 47.8% | (0.414, 0.542) |
|  | Control | 171 | 344 | 49.7% | (0.443, 0.551) |
| 5 | RTV | 121 | 258 | 46.9% | (0.407, 0.532) |
|  | Control | 182 | 354 | 51.4% | (0.461, 0.567) |
| Total | RTV | 498 | 1253 | 39.7% | (0.370, 0.425) |
|  | Control | 614 | 1507 | 40.7% | (0.382, 0.433) |

The results of the GLMM estimate that the probability of mouse infection increased among RTV and Ctrl sites over the study period; however, Ctrl sites had a higher mean probability of mouse infection from baseline compared with RTV sites for all years. The results of the model suggest that the estimated mean proportion of mice infected with *B. burgdorferi* increased by a smaller amount each year among RTV sites compared with Ctrl sites (Figure 5B; Table 6). The results of the model suggest a 7.3% (Cr-I: -2.9%, 17.5%) reduction in the proportion of mice infected with *B. burgdorferi* from baseline among RTV sites in the final year of the study compared with Ctrl sites and a 0.918 probability that the RTV resulted in a positive effect in the final year of the study.

**Table 6.** Estimated Effect on Mouse Infection from Baseline (Ctrl-RTV)

| Year | Mean Difference | Median Difference | 95% Credible Interval | Probability of Direction |
| --- | --- | --- | --- | --- |
| Baseline | - | - | - | - |
| 2 | 0.005 | 0.005 | (-0.017,0.028) | 0.673 |
| 3 | 0.049 | 0.049 | (0.002,0.096) | 0.980 |
| 4 | 0.061 | 0.061 | (-0.013,0.135) | 0.948 |
| 5 | 0.073 | 0.073 | (-0.029,0.175) | 0.918 |

A total of 1,246 mouse engorged nymphal ticks were tested for *B. burgdorferi* infection. The unadjusted observed proportion of infection prevalence among nymphal ticks collected from mice did not differ between RTV (Overall: 32.1%; Range:22.6-37.9%) and Ctrl (Overall: 32.4%; Range:28.3-34.5%) sites (Supplementary Table 2). When compared to baseline, proportions of infected engorged ticks from mice collected from RTV sites decreased every year except year 3, while Ctrl site proportions mostly increased from baseline (Figure 6A).

**Figure 6.**
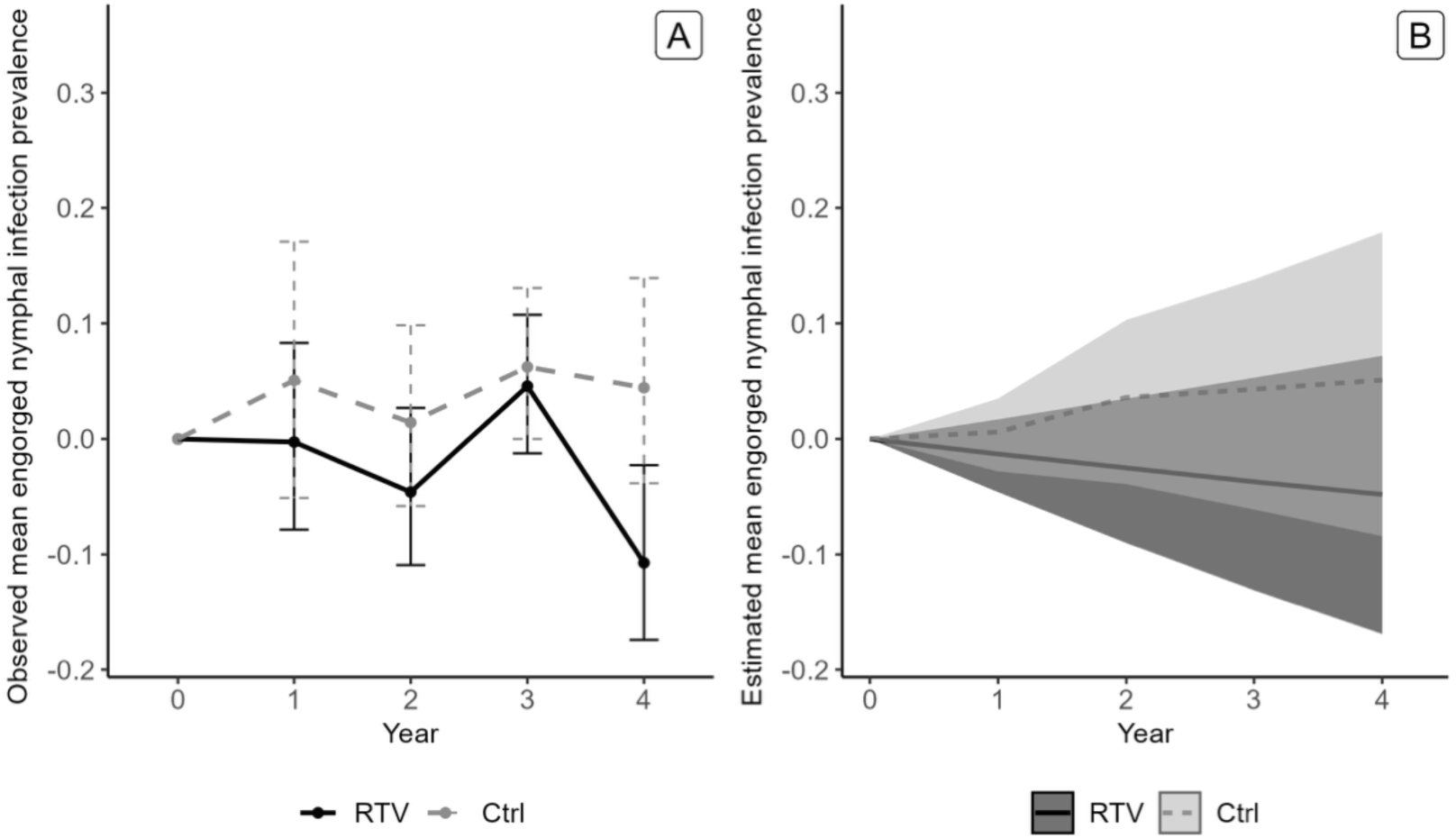
Mouse engorged nymphal tick infection prevalence with B. burgdorferi (qPCR) by year from baseline. A. Unadjusted observed proportion including 95% confidence intervals; B. Estimated mean generated from GLMM posterior samples; the shadowing represents 95% Bayesian credible intervals.

The GLMM mean estimated probability of mouse tick (engorged nymph) infection decreased for RTV from baseline to year five and Ctrl sites had a larger mean predicted probability of mouse tick infection from baseline compared to RTV sites for all years. We estimated a 9.8% (Cr-I: - 8.1%, 27.6%) reduction in the proportion of mouse tick infected with *B. burgdorferi* from baseline among RTV sites in the final year of the study compared with Ctrl sites and a 0.860 probability that the RTV resulted in a positive effect in the final year of the study (Figure 6B, Table 7).

**Table 7.** Estimated Effect on Mouse Engorged Nymphal Ticks from Baseline (Ctrl-RTV)

| Year | Mean Difference | Median Difference | 95% Credible Interval | Probability of Direction |
| --- | --- | --- | --- | --- |
| Baseline | - | - | - | - |
| 2 | 0.019 | 0.019 | (-0.026,0.063) | 0.793 |
| 3 | 0.061 | 0.061 | (-0.035,0.155) | 0.894 |
| 4 | 0.079 | 0.080 | (-0.058,0.214) | 0.875 |
| 5 | 0.098 | 0.099 | (-0.081,0.276) | 0.860 |

### Dogs

A total of 2,093 dog blood samples from 483 unique dogs were tested by PepVF ELISA. There was a very high *B. burgdorferi* exposure rate at the start of the study. In the first year, 52.1% (125/240) of dogs were seropositive for *B. burgdorferi* in August; RTV sites had 36.2% seropositivity and Ctrl sites had 73.5% seropositivity. Over the study period, 148 (19.0%) dogs seroconverted from negative *B. burgdorferi* in August to positive in the Fall with a range of 4 (2.2%) to 62 (37.8%) seroconversions among years. Among RTV sites, 71 (16.6%) of dogs seroconverted overall with a range of 3 (2.9%) to 35 (38.9%) among years. Among Ctrl sites, 77 (21.9%) of dogs seroconverted overall with a range of 1(1.4%) to 27 (36.5%) among years (Table 8). The proportions of dogs that seroconverted did not significantly differ between RTV and CTRL sites (p=0.059) and change from baseline followed a similar trend between the two groups (Figure 7A),

**Table 8.** Unadjusted observed dog Seroconversions (negative to positive) to B. burgdorferi by Treatment Group and Year.

| Year | Treatment Group | Seroconversions (negative to positive) | Total | Proportion | 95% CI |
| --- | --- | --- | --- | --- | --- |
| Baseline | RTV | 3 | 81 | 3.7% | (0.008, 0.104) |
|  | Control | 3 | 26 | 11.5% | (0.024, 0.302) |
| 2 | RTV | 3 | 104 | 2.9% | (0.006, 0.082) |
|  | Control | 1 | 74 | 1.4% | (0.000, 0.073) |
| 3 | RTV | 35 | 90 | 38.9% | (0.288, 0.497) |
|  | Control | 27 | 74 | 36.5% | (0.256, 0.485) |
| 4 | RTV | 18 | 83 | 21.7% | (0.134, 0.321) |
|  | Control | 22 | 72 | 30.6% | (0.202, 0.425) |
| 5 | RTV | 12 | 71 | 16.9% | (0.090, 0.277) |
|  | Control | 24 | 106 | 22.6% | (0.151, 0.318) |
| Total | RTV | 71 | 429 | 16.6% | (0.132, 0.204) |
|  | Control | 77 | 352 | 21.9% | (0.177, 0.266) |

**Figure 7.**
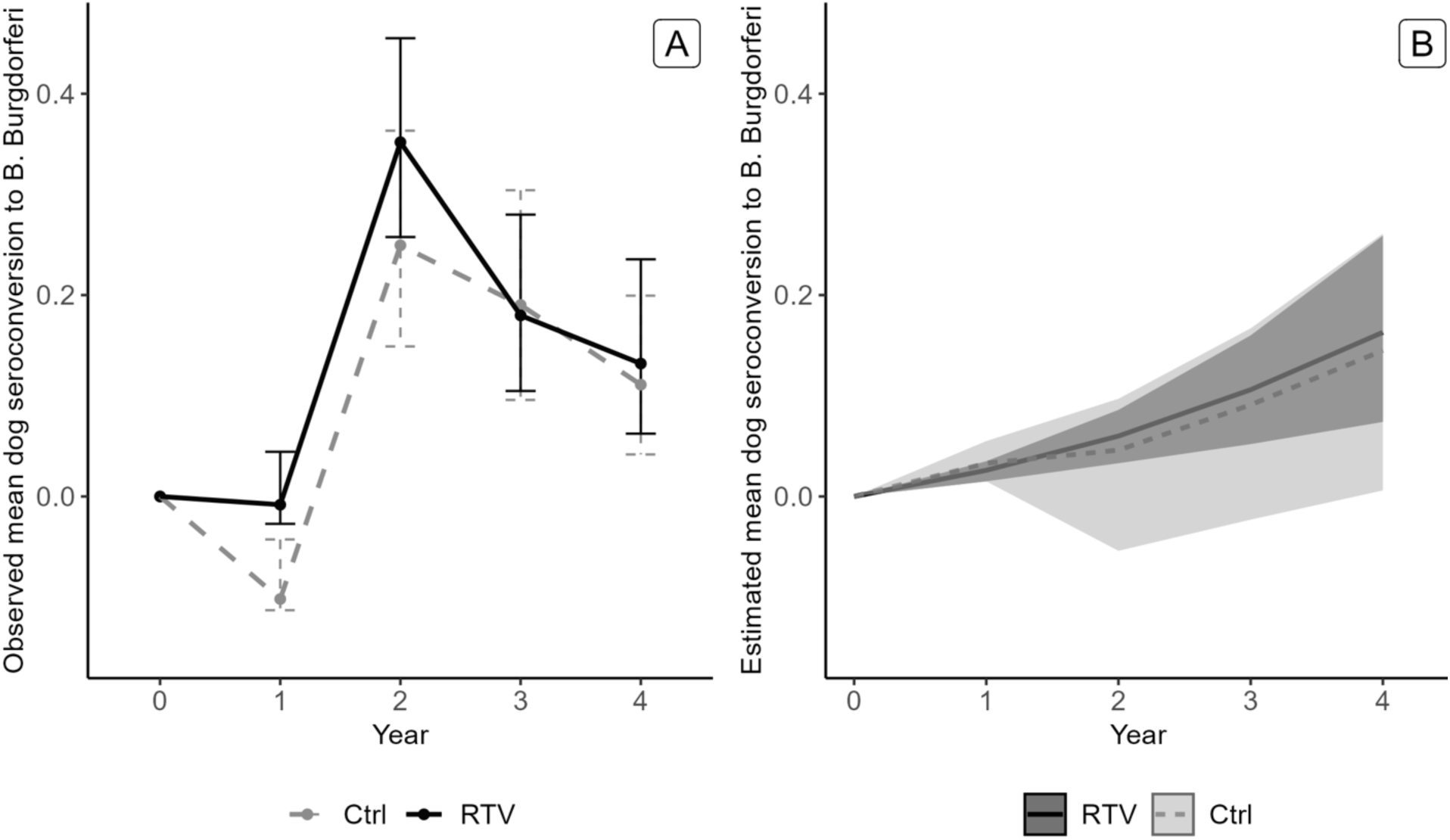
Dog seroconversion to B. burgdorferi (negative to positive) rate by year. A. Unadjusted observed proportion including 95% confidence intervals; B. Estimated mean generated from GLMM posterior samples; the shadowing represents 95% Bayesian credible intervals.

The results of the GLMM suggested that the probability of infection increased over the study period among dogs at RTV and Ctrl sites (Figure 7B; Table 9). From the baseline to the final year of the study, the estimated mean probability of infection increased among dogs at RTV sites by 1.8% (Cr-I: -0.180,0.129) compared to Ctrl sites. The probability that the RTV resulted in a positive effect on dog seroconversion was 0.58 for the last year of the study compared to baseline.

**Table 9.** Estimated Effect on Dog Seroconversion (negative to positive) to B. burgdorferi from Baseline (Ctrl-RTV)

| Year | Mean Difference | Median Difference | 95% Credible Interval | Probability of Direction |
| --- | --- | --- | --- | --- |
| Baseline | - | - | - | - |
| 2 | 0.007 | 0.007 | (-0.013,0.031) | 0.748 |
| 3 | -0.015 | -0.008 | (-0.115,0.046) | 0.604 |
| 4 | -0.015 | -0.011 | (-0.136,0.080) | 0.586 |
| 5 | -0.018 | -0.015 | (-0.180,0.129) | 0.580 |

Furthermore, a total of 166 adult *Ixodes scapularis* ticks were collected from dogs over the study period; 39 ticks were collected from dogs residing near RTV sites and 44 ticks were collected from dogs residing near Ctrl sites. Approximately 18% of *I. scapularis* ticks collected from dogs were infected with *B. burgdorferi.* A slightly larger proportion of ticks from Ctrl sites were infected (20.5%) compared to RTV sites (15.4%); however, the difference was not statistically significant (Supplementary Table 3).

A summary of the unadjusted observed non-normalized data is shown for flat nymphal *B. burgdorferi* infection prevalence, mouse engorged nymphal *B. burgdorferi* infection prevalence, mouse ear *B. burgdorferi* infection prevalence, mouse OspA seroprevalence, and dog seroconversion to *B. burgdorferi* among Ctrl and RTV sites (Supplementary Figure 1).

## Discussion

We conducted a 5-year double blinded placebo-controlled study of a reservoir targeted vaccine (RTV) and were able to demonstrate impacts across several different components of the *B. burgdorferi* enzootic cycle. This study is the first to evaluate deployment of RTV in multiple field sites for 5 consecutive years and to utilize the dog as a proxy for prediction of human Lyme disease risk. Our results show strong evidence of increased OspA seroprevalence among mice on RTV sites compared to control (Ctrl) sites indicating that the vaccine was broadly consumed by *Peromyscus*. Probability estimates from the GLMM model, which captures site-to-site variation as well as overall group effects, indicated strong evidence of protective levels of antibodies to OspA among mice captured in RTV field sites (10.5%) and a 15.4% reduction in NIP in RTV treated sites by the 5^th^ year of the study. Furthermore, mouse ear infection prevalence did not increase in RTV sites as much as it increased in controls (difference 7.3%) and mouse engorged nymphal tick prevalence was reduced by 9.8%. Lastly, our current study design did not find evidence of an impact on dog seropositivity, raising questions about the usefulness of heavily exposed cohorts of working dogs as proxy indicators for human Lyme disease risk.

Our findings are consistent with other studies that have evaluated reservoir-targeted oral OspA vaccines. Richer et al. found that NIP decreased by 76% after 5 years of RTV deployment on one of the four New York treatment sites (compared to a 94% increase on control sites) and site-specific effects led to a significant decrease in NIP by year three in additional sites, which was linked to increased anti-OspA seroprevalence in *Peromyscus*^13^. These findings are similar to the unadjusted observed findings in the current study of a 42.6% decrease in NIP (22% to 12.7%) among treatment sites compared to a 148.3% increase in control sites (6.5% to 16.2%) which is associated to increased OspA seroprevalence. Our stringent probability estimates from the GLMM indicated strong evidence of a 15.4% reduction in NIP in RTV treated sites by the 5^th^ year of this study.

The GLMM estimated a 10.5% increase of protective anti-OspA antibody over the 5-year study. However, comparing the unadjusted proportion of protective OspA antibodies among mice captured in RTV field sites over 5-years (15.3%) was slightly lower in the current study than OspA seroprevalence measured in the RTV plots that produced significant NIP differences the first field study over the term of the study (21-33%)^13^. Although we delivered 1/3 of the RTV in Sherman traps as a backup to carousel malfunction, we delivered most (2/3) of the oral RTV via automatic carousels in realistic field conditions. The carousel was designed for deployment in peridomestic settings and each carousel provides vaccine for a 1/2-acre radius. Thus, 8 carousels should have been sufficient for each 1.1 hectare field site for optimal delivery. Although protective OspA antibodies are transferred from *Peromyscus* dams to pups via lactation^17^, mouse population turnover may have impacted OspA seroprevalence as mice do not usually live more than one year in natural ecosystems (in our study only 10% of mice were recaptured the following year). Thus, significant OspA seropositivity is not expected to carry over into the following year. We provided bait vaccine almost all year around as we have shown that prolonged vaccination of *Peromyscus* maintained higher levels of anti-OspA antibody ^20^. Another factor to consider is that in the current study we used the inactivated version of *E. coli*-OspA which was the composition under consideration for licensing approval by the USDA, whereas in the first study we used a live *E. coli*-OspA vaccine. Ultimately, these factors may have contributed to reduced number of mice with OspA titers above the correlate of protection threshold. The presence of other rodents assessing the carousels would have not impacted OspA seroprevalence in mice, as the latter depends on availability of bait, which was available throughout most of the year. We did trap voles and chipmuncks in some sites. Consumption of baited vaccine by other species may have contributed to the reductions in NIP observed, although we did not assess that factor specificaly. One factor that could have affected consumption of OspA-RTV could be pallatability of the bait which was not assessed in this study. Thus, strategies to improve RTV coverage to all rodent reservoir hosts, as well as strategies that increase the titer of OspA antibody in *Peromyscus* would likely further reduce NIP.

Although the GLMM mean probability estimate of *B. burgdorferi* infection in mice increased among RTV and Ctrl sites over the study period, it was substantially lower in RTV than in Ctrl sites with a measurable difference of 7.3%. In addition, the GLMM effect size shows a 9.8% reduction in mouse engorged nymph infection prevalence. A two-year evaluation of RTV conducted in Connecticut resulted in reduced proportions of *Peromyscus* with *B. burgdorferi* infection on RTV sites as compared to control sites as well as reduced infection prevalence among larval *Ixodes* feeding on mice ^14^, much in line with the carousel based broader dispersion of RTV and findings of the current trial.

In absolute terms, the GLMM effect size (15.4% reductions in NIP, 9.8% reduction in mouse engorged nymph infection prevalence and 7.3% lower mouse ear infection prevalence) is attributable to the proportions of protective levels of anti-OspA antibody (10.5%) in *Peromyscus* in RTV sites. Our data strongly suggests that anti-OspA antibody present in the blood of RTV-vaccinated/infected *P. leucopus* reduce *B. burgdorferi* infection prevalence in the larval stadial stage when they become active in the summer (1^st^ year of the enzootic cycle) before they molt into nymphal ticks that will quest the following spring (qNIP) without substantially reducing infection prevalence of the mouse. Thus, OspA-RTV reduces transstadial transmission of *B. burgdorferi*, larva to nymph and likely nymph to adult. The following findings from other studies help substantiate our conclusion. OspA is upregulated by *B. burgdorferi* after it enters naïve larval ticks and OspA is also detected on the surface of *B. burgdorferi* in flat unfed nymphal ticks^18,19^ showing that OspA is essential for *B. burgdorferi* colonization of the tick vector.

Furthermore, antibody to OspA induced in *Mus* and *Peromyscus* after OspA-immunization has been shown to block transmission of *B. burgdorferi* within the tick^18, 20^. It has also been shown that OspA vaccination of previously infected mice including *Peromyscus* decreased acquisition of *B. burgdorferi* to larval *I. scapularis* ticks^19^ ^20^ and that this effect was linked to increased vaccination doses, which increases titers of antibodies to OspA ^20^. However, OspA antibodies circulating in the blood of mice do not reduce *B. burgdorferi* load in infected mice^21^ because *B. burgdorferi* downregulates OspA in the nymphal tick midgut prior to transmission to the vertebrate host^22,23^.

Potential differences among sites could have impacted results through multiple mechanisms. At baseline, due to random chance, the RTV treatment sites had substantially higher NIP than control sites, indicating considerable variability among sites. Another factor to consider is that NIP is typically lower (∼16%) in Maryland compared to states in the Northeast^24^. Other ecological factors affecting the enzootic cycle such as higher proportions of other hosts in the sites (shrews and chipmunks) which have been shown to play important roles as reservoir hosts^19,25,26^, fluctuating abundance of mice and seasonal mouse demography^27–29^ could have obscured our ability to distinguish RTV-effects in mice. In addition, we must account for unpredictable variables that cannot be controlled. For example, the study was launched during the 2020 Covid shutdown, a tornado hit several field sites in 2023 that displaced traps and carousels and possibly the reservoir population. Although we regularly checked carousels throughout the summer, there were times when carousels were not operating due to damage from natural conditions. Situations included damage or wear from rodent gnawing and environmental conditions (e.g., water and storm damage) and displacement of carousels due to animal interference, mud slides, or tree damage. Certain sites were more prone to damage due to land characteristics and local ecology.

We found no evidence of a difference in seroconversion (negative to positive) between dogs living on RTV sites compared to those living on Ctrl sites. There are several potential explanations for this finding. First, it is possible that the impact on NIP was not substantial enough to impact Lyme disease risk in dogs. Due to the high correlation between risk of an incidental host contracting LD and nymphal infection prevalence ^30–35^, based on mathematical modeling, even a ∼12% decrease in nymphal infection prevalence in initial years of deployment could lead to significant reductions in human incidence of LD ^36^.The relationship between nymphal infection prevalence and Lyme disease risk is still not well defined, and it is unclear the reductions in NIP that must be achieved for meaningful epidemiological impact in human disease outcomes^37^, or in a highly exposed population like hunting dogs. Second, some evidence suggests that dogs are more likely to acquire *B. burgdorferi* from adult ticks compared to nymphal ticks^38^, however, adult tick season often peaks in November in Maryland^39^. Therefore, it is possible that dogs infected between August and November may not have had sufficient time to develop detectable antibodies^38^. Finally, despite rather limited exposures outside of the experimental fields, dogs were exposed to areas outside of field plots from September-February and may have had exposure during these times leading to null differences in canine serology between the RTV and Ctrl groups. This of course would be a concern for people using similar techniques to prevent exposure at home but then could easily get exposed to an infected tick out on a hike at a nearby state park. As a part of this trial, we asked that other interventions not be used if not part of their current protocols and this was observed during the five years of study. This may have put dogs at greater risk when hunting away from the experimental fields. All interventions should be considered, like use of appropriate acaracides with an ecological RTV, as a package to prevent overall LD risk.

Our study was subject to some limitations. First, we observed substantial site-to-site and year-to-year variation in RTV effect, indicating that the impact of RTV may be inconsistent at larger geographic scales and varying landscapes. The models were chosen to be able to account for overall differences between sites (using random intercepts), to account for random variation (via the binomial likelihood), and to allow estimation of average changes over time within sites and within groups of sites (via the time-group interaction). Nevertheless, the generalizability of the findings is necessarily limited by the characteristics of the sites under study. Second, control field site 1.1 was replaced after the second year of the study due to distinct focal ecology predominantly dominated by guineafowl which are known to be avid consumers of ticks. Given this challenge, field site 1.2 replaced the initial site beginning in year three. Therefore, we were unable to observe trends across the full 5-year period for these locations. Third, potential differences among sites could have impacted results through multiple mechanisms. At baseline, due to random chance, the treatment sites had substantially higher NIP than control sites, indicating considerable variability among sites. Given this variability, other unobserved site-specific differences could have obscured our ability to distinguish effects. Fourth, dragging practices also varied slightly by site owing to a handful of factors. First, sites had varying landscape types with some more suitable to drag sampling while others were comprised of denser vegetation, complicating dragging efforts. As a result, drag-sampling was concentrated to areas without barriers that impeded dragging and dragging frequency varied by site. Finally, the study design made estimating DIN unfeasible. For that measure we would need to have enacted a drag protocol to cover a full hectare (100 m^2^ and ideally more - 750 m^2^), as recommended by the Centers for Disease Control and Prevention. Instead dragging was carried out 0-20 meters from the trap lines until a minimum number of ticks was found for estimating NIP.

Our study indicates that use of an OspA-based vaccine delivered orally by inactivated cells of *E. coli* (RTV) was associated with increased proportions of mice with anti-OspA antibodies in RTV treated sites (GLMM estimate 10.5%), as well as with reductions in the proportion of infected questing nymphal ticks (GLMM estimate 15.4%). Our data strongly suggests that anti-OspA antibody in infected-vaccinated *P. leucopus* reduce larval infection prevalence when mice consume RTV in the first year of the enzootic cycle which we measure as reductions of nymphal infection prevalence the following spring without affecting mouse infection prevalence substantially. This further suggests that OspA-RTV reduces transstadial transmission of *B. burgdorferi*. Methods to improve broadcast delivery of RTV to additional hosts and to increase titers of anti-OspA antibody in reservoir hosts are likely to further reduce NIP. RTV could be an important tool in a set of interventions for Lyme disease prevention on both residential and commercial property. Our study arm designed to evaluate potential impact of RTV treatment to incidence of Lyme disease in another incidental host of this spirochete was inconclusive. Further study is needed to determine how RTV impacts human Lyme disease incidence as well as longer-term use of RTV in the ecotone.

## Methods

### Guidelines

All guidelines were followed in accordance with protocols approved by the University of Iowa Institutional Animal Care and Use Committee and the University of Tennessee Health Science Center.

### Field Sites

The study was conducted in four Maryland counties (Baltimore, Harford, Howard, and Montgomery) at seven privately owned 1.1-hectare (ha) field sites (Figure 1) over a 5-year period (2020-2024). Each field site was adjacent to a hunting dog kennel and field sites were regularly used as exercise grounds for the hunting dogs during March through August. Dogs continued to be exercised in these areas in the remaining months of the year (September-February) but were engaged in hunting away from field sites 2-3 times a week. Ecological features of field sites are described in detail elsewhere^40^. Briefly, sites were rural to suburban peridomestic and contained potential tick habitat. One field site was replaced after the second year of the study due to inability to collect sufficient numbers of *Ixodes scapularis* ticks during dragging activities. Field sites were paired by geographic location with sites located within 2.5- 11.5 km of each other. Block randomization was conducted, wherein for each of the three geographic locations, one site was randomized to RTV and the other was assigned to Ctrl^41^. All field team members were blinded to treatment assignments for the entirety of the study, members of the Gomes-Solecki laboratory were blinded during assay performance, and the statistical team was unblinded over the entirety of the study.

### Bait preparation and quality control

RTV bait was prepared by the Gomes-Solecki lab according to previously published protocols^14,42,43^. Briefly, a culture of *Escherichia coli* expressing *B. burgdorferi* OspA B31 was chemically inactivated with beta-propiolactone (BPL) following a protocol approved by the USDA-CVB to produce an inactive whole-cell vehicle that was used to coat non-nutritious LabDiet mouse pellets (Purina, Largo, FL) containing ∼ 6 µg of OspA per pellet. Non-nutritious pellets were used to prevent unintended increases in mouse populations in the field sites. Inactivation of *E. coli* was confirmed by culture of BPL-treated and untreated *Escherichia coli* expressing *B. burgdorferi* OspA in TBY media containing 50 µg/mL kanamycin, which was measured by spectrophotometric reading at OD_600_. Uncoated pellets were used for placebo (Ctrl) bait.

The RTV bait was subjected to quality control analysis as follows: groups of C3H-HeN mice (n=3 to 5, per lot) received oral bait vaccinations following our standard schedule of immunization comprised of a prime over 2 weeks, a 1-week rest and a 2-week boost. Each mouse received a total of 20 bait pellets per week of vaccination. Low volumes of blood (<50 µl) were collected by tail nick or from the submandibular vein on days D0, D17, D38, D59, D62 and D72 for determination of anti-OspA antibody by ELISA. Two weeks after the last vaccine dose, challenge was done with laboratory produced *Ixodes scapularis* carrying > than 10 strains of *B. burgdorferi* and mice were euthanized 3 weeks after the last day of challenge (D72) for collection of blood for quantification of anti-*B. burgdorferi* antibodies by pepVF ELISA^44,45^, as well as heart tissue which was processed for determination of *B. burgdorferi* DNA by *flaB* quantitative PCR (qPCR). Blood collections and challenges were performed without anesthesia by experienced personnel. Euthanasia was performed in mice anesthetized with isoflurane 3-5% followed by exsanguination and thoracotomy to collect tissues.

RTV and Ctrl bait samples from each production lot were stored in the lab throughout the year and subjected to quality control protocols. Pails containing 20,000 dry bait pellets (3 pails of RTV and 3 pails of Ctrl) were blinded by color coding and shipped to the field sites in Maryland each year, where they were stored in an outdoor sheltered location and exposed to variable temperatures for up to a year. Bait pellets stored for up to a year were validated for use after passing our vaccine efficacy quality control analysis.

### Carousel filling and placement

Bait was dispensed by automatic carousels (US BIOLOGIC, Inc, Supplementary Figure 2), spaced approximately 50 m apart along mouse trapping lines at study sites where the ecotone was favorable to the presence of small rodents. Carousels were placed 50-100m from kennels and any domiciles in wooded areas at least 5m from the forest edge. Carousels contained 6 compartments holding ∼100 pellets and rotated every 10 days, thus providing bait for a 60-day period. Eight (8) carousels were deployed at each field site (1.1 ha). Carousels were refilled and checked for proper placement and function in March, May, July, August, and November of each study year to ensure that bait was available to mice from March to January of the following year. Overall, roughly 2/3 of the bait vaccine (RTV) shipped to the field sites was deployed in automatic carousels and 1/3 was deployed manually in Sherman traps as a backup to ensure sufficient bait availability for adequate protection in the RTV group.

### Mice

During May through July of each study year, live Sherman traps (3×3×10”, Tallahassee, FL) were deployed at each study site on alternating weeks for 4 days per week; each trap was set with three bait pellets (RTV or placebo) with the goal of capturing *Peromyscus* spp mice. Traps were set in the evenings and checked in the mornings by 11 am. Trap lines were replaced each March via survey tool, measuring 5m from the forest edge and 15 m between lines. Each year trapping efficiency was assessed and optimized for mouse capture at each of the sites. Because of this, by the end of the trial the number of traps varied by site with a minimum of 19 (very high trap capture rates) and a maximum of 42 (lower per trap capture rates). A map of the layout of trap lines relative to the property is shown (Supplementary Figure 3). Captured mice were manually restrained, tagged with a unique identifier, and approximately 20-50 µl blood sample and 2 mm diameter ear punch sample were collected once per trapping week at maximum with the goal of collecting samples from at least 30 unique mice per site each field season. Information regarding age, sex, recapture, and proportion of bait consumed were recorded upon each mouse capture as detailed previously^40,44^. When present, nymphal and adult ticks were collected from captured mice. No field mice were anesthetized or euthanized during the study. Mice were released into the field at the trapping location after sample collection was completed.

### Drag Ticks

Tick dragging activities were conducted in peak nymphal questing season during May through July of each study year on alternating weeks with the goal of collecting at least 30 flat unfed nymphal *Ixodes scapularis* ticks per site. We started Mid-May each year based on phenology of *Ixodes* after conferring with Dr. Ellen Stromdahl ^39,46^, see Suppl. Fig. 3. Drag sites were on or adjacent to trap lines, optimizing sites based on leaf litter and other ecological factors for tick retrieval. When at least 30 flat unfed nymphal *Ixodes* ticks were collected at a specific site, efforts were shifted to remaining sites for which tick collection goals had yet to be met. A flannel drag cloth approximately 1m^2^ in size was dragged along areas of tick habitat within a 1-acre radius of carousels. Drag cloths were checked for ticks every 15 m of distance dragged. Ticks collected by dragging or found crawling on field staff engaged in dragging were collected for identification and testing.

### Dogs

To assess canine exposure to *B. burgdorferi*, which includes some nymphal exposures, but is thought to be primarily via adult ticks^38^ (Sup. Figure 3), hunting dogs living in kennels adjacent to the field study plots were physically examined annually in August and information regarding lymph node size and physical condition were recorded. Dog breeds housed by the kennels varied across sites and included foxhounds (4 sites), beagles (1 site), and basset hounds (2 sites). Exclusion criteria included hounds and beagles younger than 3 months or older than 4 years. This was because older dogs are more likely to have lingering *B. burgdorferi* antibodies^15^ and younger dogs are less likely to have tick exposure. During physical examination, blood was collected for Lyme disease serology testing and any adult *Ixodes scapularis* ticks noted on examined dogs were collected beginning in the second year of the study. Kennel masters were also provided with tick collection supplies to collect ticks found attached to dogs during the study period. All collected ticks were tested by qPCR for presence of *B. burgdorferi flaB*. Dog serum was collected for determination of antibodies against *B. burgdorferi*. Dogs who tested negative in August were examined and tested again in the fall (October or November) to determine if seroconversion had occurred. Dogs who seroconverted from negative to positive between August and November were considered to have had an incident *B. burgdorferi* infection. Samples with missing identification numbers were excluded from analysis. No dogs were anesthetized or euthanized during the study. Dogs did not receive previous vaccination against Lyme disease and were not treated with anti-tick medications. In Supplementary Figure 4 we show the study design with spatial extent for dragging of questing nymphs, deployment of baits, and sampling of mice and dogs.

### Tick Identification and B. burgdorferi diagnostic methods

Trained staff examined all ticks collected from dogs, mice, and dragging activities using a dissecting microscope to assess species, life stage, and engorgement status. Flagellin B (*flaB*) qPCR was used to determine presence of *B. burgdorferi* spirochetes in *Ixodes scapularis* ticks and mouse ear tissue; a cycle threshold of 50 was used to determine presence of *B. burgdorferi*. ELISA was used to evaluate anti-OspA antibody levels in mice according to previously described protocols^15^; protective OspA seropositivity was defined as optical density (OD) at 450 ≥ 0.80 as an approximate correlate of protection ^43^. PepVF ELISA was used to determine presence of *B. burgdorferi* antibodies in mouse and dog serum samples according to described protocols^45,47^.

### Statistical Analysis

Unadjusted aggregate nymphal tick infection prevalences were calculated for each year for RTV and Ctrl sites. For analysis of optical density (OD) data on anti-OspA seroprevalence, unpaired t tests with Welch’s correction were used to compare RTV with Ctrl sites. In addition, the proportion of mouse blood samples that tested positive for anti-OspA antibodies above the predicted correlate of protection threshold^43^ were compared between RTV and Ctrl control groups. Likewise, the proportion of ear samples and flat and engorged *Ixodes scapularis* ticks (nymphs from drag and mice, and adult ticks from dogs,) that tested positive for *B. burgdorferi* were compared between RTV and control groups.

*B. burgdorferi* seroconversion rates among dogs were calculated for each year to determine the proportion of dogs testing negative in August who developed antibodies against *B. burgdorferi* at the time of fall testing in October/November. These seroconversion rates were compared between RTV and Ctrl groups. Comparisons of proportions between RTV and Ctrl sites were conducted using chi-squared tests or Fisher’s Exact tests where appropriate (α = 0.05). Analysis of difference in differences of proportions [(year 5 RTV – year 1 RTV) – (year 5 Ctrl – year 1 Ctrl)] was done using a normal approximation. Marginal confidence intervals were computed using exact binomial intervals^48^.

### Model

We used a Bayesian Generalized Linear Mixed Model (GLMM) with a logit-link function^49^ to estimate the effects of RTV. The model included variables for treatment, year, and the interaction between treatment and year as fixed effects. Random intercepts for the sites were incorporated to account for natural variations that may influence anti-OspA antibody levels and *B. burgdorferi* infection prevalence. This selection of terms allows each site to have a separate linear trajectory for the outcomes over the study period, while retaining our ability to aggregate information from the treatment and control sites to obtain effect estimates. The use of a Bayesian framework for the primary model comparison eliminates the need to rely on binary decision thresholds in a relatively small study, allowing us to directly report the remaining posterior uncertainty after having observed the data.

The statistical analysis was performed using the rstan package in R^50^ to obtain posterior samples of model parameters and derive quantities through Markov chain Monte Carlo (MCMC) techniques. We specified independent weakly informative normal priors with mean 0 and variance 10 for the fixed effects, and a multivariate normal distribution with Cholesky parameterization for the random effects with mean 0. The posterior results are based on a set of (3) MCMC chains of 20000 iterations on each chain, after the burn-in period. To assess the level of convergence of the parameters, the Gelman-Rubin diagnostic was computed using the Coda package in R^51^.

Posterior samples for model parameters were summarized by calculating posterior means, medians, probability of direction (the estimated probability that the difference is positive or negative), and lower and upper values of the 95% credible intervals (Cr-I) in each group for mouse infection with *B. burgdorferi*, mouse protective OspA seroprevalence, dog infection, tick infection with *B. burgdorferi* (flat nymphal tick and mouse engorged nymphal tick) and dog seroconversion to *B. burgdorferi*. Serologic evidence of exposure to *B. burgdorferi* in mouse was not included in the models. Posterior differences (Ctrl vs. RTV) between posterior probabilities of prevalence/incidence were calculated to assess the effect of RTV. These measures were calculated on a normalized scale (additive change relative to baseline) for all estimates except for OspA seroprevalence estimates. Probabilities near 0.5 indicate no evidence either way, while probabilities near 1 or zero indicate strong evidence of the directional effect. We consider probabilities between 0.2 to 0.8 to correspond to equivocal or weak evidence, with 0.8-0.9 (0.1-0.2) representing strong evidence, and greater than 0.9 (or < 0.1) representing very strong evidence in favor of a directional effect. The study was powered based on an estimate of a minimum of 30 flat, nymphal ticks per site per year, estimating a minimum expected effect size of 15% decrease in NIP for a corresponding power of > 90% at a type-1 error rate of 90%. Mouse targets were chosen to match. Finally, Bayesian GLMMs were selected for the primary study comparisons due to their lack of reliance on binary decision rules; instead of a binary declaration of significance, the probability of positive/negative effects can be directly computed (probability of direction).

## Data availability

The processed experimental data that support the findings of this study are available in GitHub with the identifier https://github.com/FerneyH/RTV

## Acknowledgements

We thank Shahriar Motazedian, Kamalika Samanta, Caroline Solecki, Chunyan Li, Aseel Alamri, Molly Solecki, Maria Sanches Pacheco and Olifan Abil for their technical contributions to this study as well as Dr. Rachel Blakey, Evelyn Cochran, and Dan O’Toole for their continued support throughout the entirety of the study. We would like to thank US BIOLOGIC Inc and Luciana Richer for technical support regarding production of the reservoir targeted vaccine. We would also like to thank the collaborating hound groups without whom this work would not be possible. This research received funding from the National Institutes of Health, National Institute of Allergy and Infectious Diseases grant numbers R01AI139267 (MGS, CP, GB), R01AI175417 (MGS), R44AI167605 (MGS).

## Author Contributions

A. M. S. data analysis and writing the manuscript

F. H-C and F. P-R. data analysis, writing part of the manuscript and created figures

K. A. and J. P. field team coordination, sample collection and processing

M.W. sample collection and processing, data analysis and writing part of the manuscript

G. J., J.F.A, S. K., N.N. vaccine production and shipping logistics, performed laboratory assays and data analysis

T. B., E. K., K. M. data collection, cleaning and analysis

P. W. and M. F. T. samples and data collection

G. B. experimental design, funding, data analysis, writing part and edited the manuscript

C. P. and M. G-S. are responsible for the concept, experimental design, funding, data analysis and edited the final version of the manuscript

All authors reviewed the manuscript.

