## Supplemental Material for "A double-blinded, placebo-controlled field trial of an OspA-based oral reservoir targeted vaccine against *Borrelia burgdorferi*"

Maryland Field Team: Briana Bowen, Kaleigh Conroy, Eli Davis, Tanner Davis, Joseph Ferguson, Lexi Frank, Victoria Kamilar, David Marquez, Carly Martin, Emily Knight, Rachel S. Krizek, Chelsi Preuc, Alec Rutherford, Phurchhoki Sherpa, Reegan Sturgeon, Derek Thorne, Jay Xiao

#co-last author/MPI team

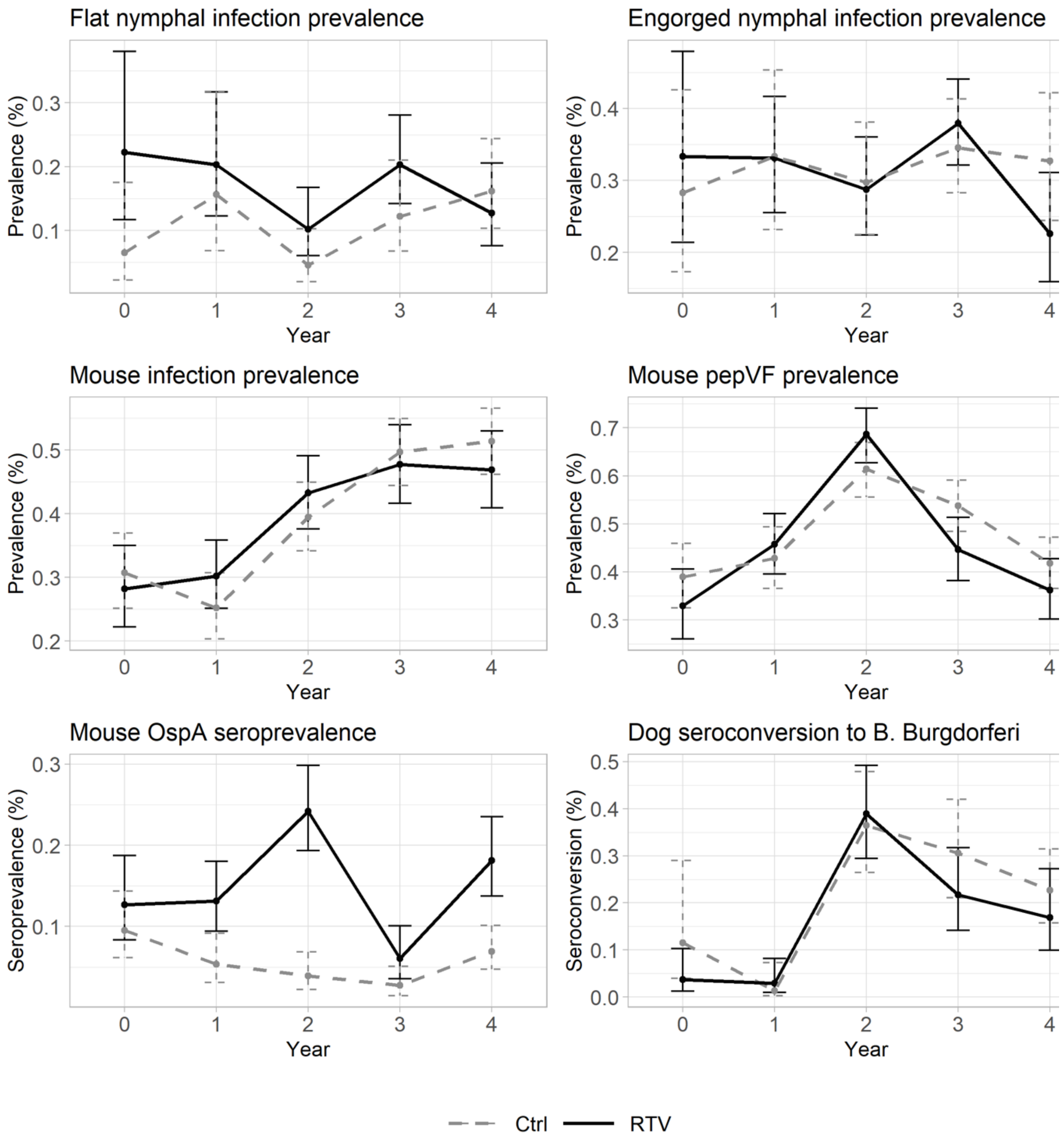


***Supplementary Figure 1.*** Summary of unadjusted observed non-normalized (+/- 95% confidence intervals) flat nymphal Bb infection prevalence, mouse engorged nymphal Bb infection prevalence, mouse ear Bb infection prevalence, mouse OspA seroprevalence, and dog seroconversion to Bb among Ctrl and RTV sites. Legend: Ctrl, control sites treated with non-vaccine bait; RTV, bait-vaccine treated sites; Bb, *Borrelia burgdorferi;* pepVF, antibody to B. burgdorferi VlsE and FlaB.

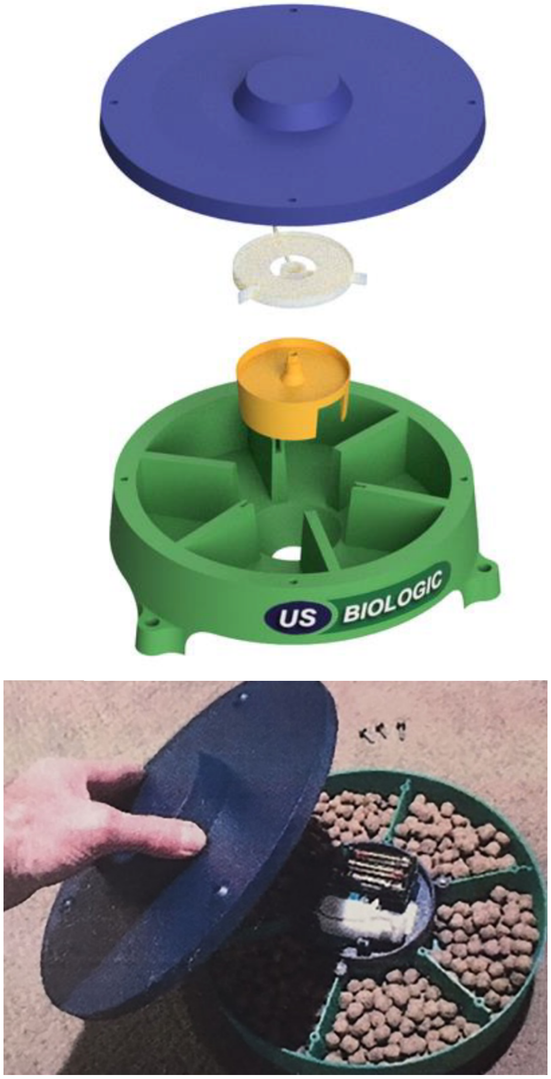


**Supplementary Figure 2.** Example of a carousel device used to distribute the RTV and Ctrl in field sites.

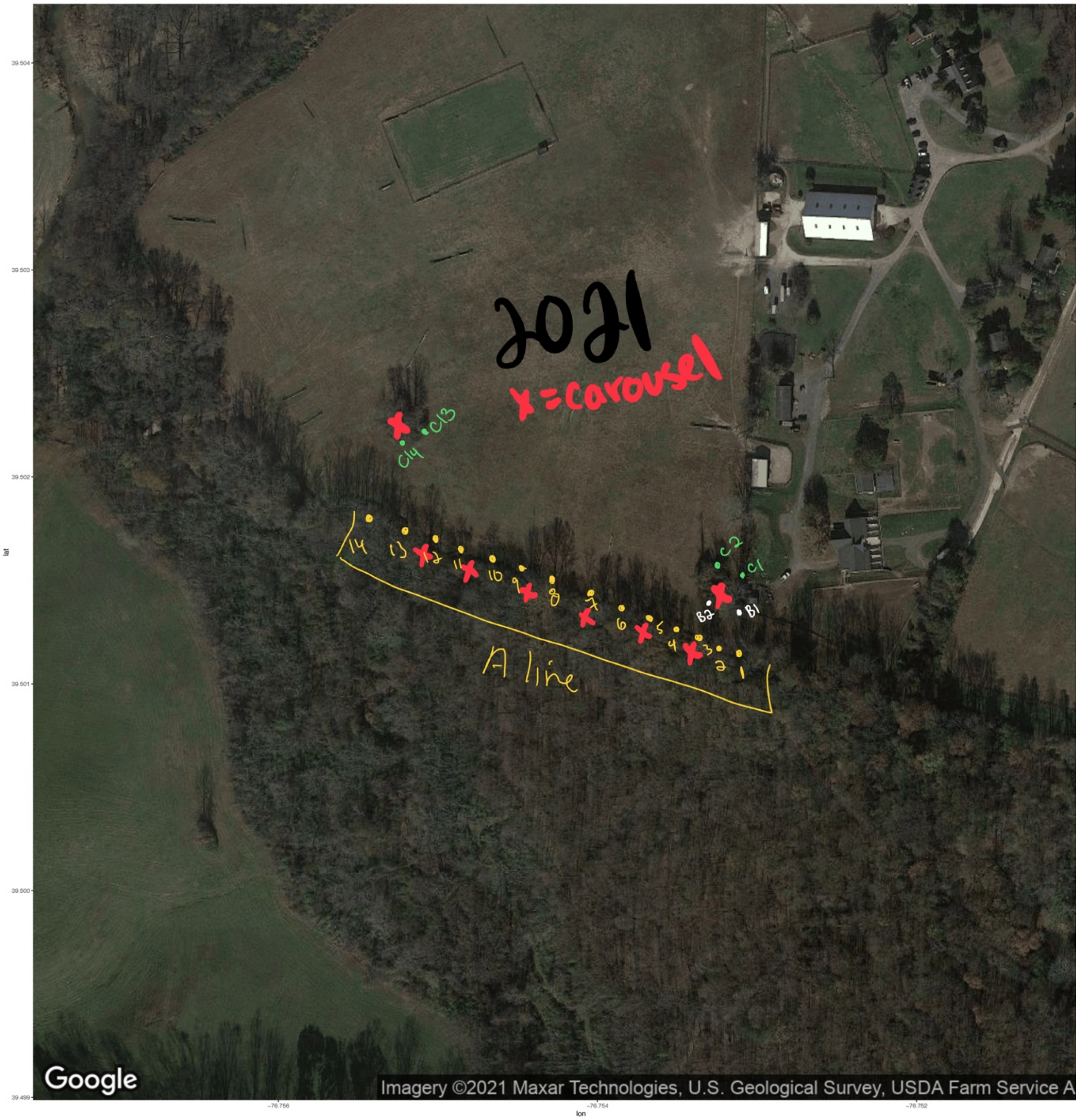


**Supplementary Figure 3.** A map of the layout of trap lines relative to the property.

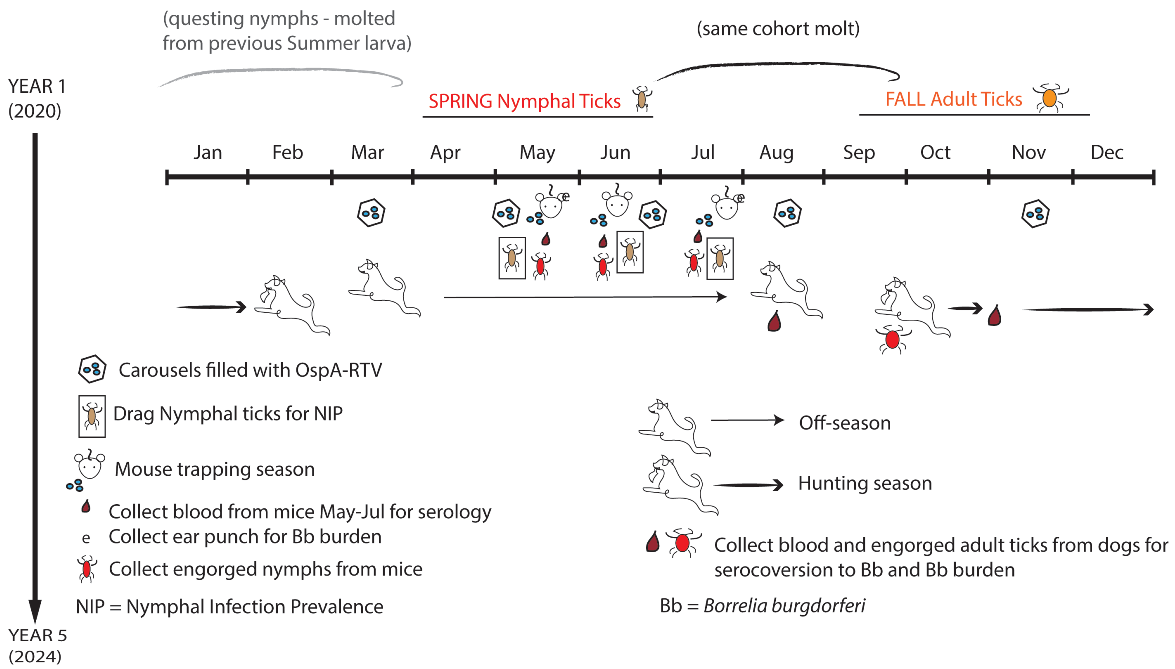


***Supplementary Figure 4.*** Study design with spatial extent for dragging of questing nymphs, deployment of bait and sampling of mice and dogs.

**SUPPLEMENTARY TABLES**

***Supplementary Table 1.*** Unadjusted observed prevalence of PepVF antibodies Among Mice by Treatment Group and Year

| Year | Treatment Group | PepVF Positive | Total Mice Tested | Proportion |
|---|---|---|---|---|
| Baseline | RTV | 52 | 158 | 32.9 |
| | Control | 78 | 200 | 39.0 |
| 2 | RTV | 108 | 236 | 45.8 |
| | Control | 96 | 224 | 42.9 |
| 3 | RTV | 173 | 252 | 68.7 |
| | Control | 172 | 280 | 61.4 |
| 4 | RTV | 96 | 215 | 44.7 |
| | Control | 178 | 331 | 53.8 |
| 5 | RTV | 80 | 221 | 36.2 |
| | Control | 135 | 323 | 41.8 |
| Total | RTV | 509 | 1,082 | 47.0 |
| | Control | 659 | 1,358 | 48.5 |

***Supplementary Table 2.*** Unadjusted observed infection Prevalence Among Mouse Engorged Nymphal Ixodes Ticks by Treatment Group and Year

| **Year** | **Treatment Group** | **FlaB Positive** | **Total Ticks Tested** | **Proportion** |
|---|---|---|---|---|
| Baseline | TBRTV | 15 | 45 | 33.3% |
| | Control | 13 | 46 | 28.3% |
| 1 | TBRTV | 42 | 127 | 33.1% |
| | Control | 22 | 66 | 33.3% |
| 2 | TBRTV | 48 | 167 | 28.7% |
| | Control | 38 | 128 | 29.7% |
| 3 | TBRTV | 94 | 248 | 37.9% |
| | Control | 69 | 200 | 34.5% |
| 4 | TBRTV | 26 | 115 | 22.6% |
| | Control | 34 | 104 | 32.7% |
| Total | TBRTV | 225 | 702 | 32.1% |
| | Control | 176 | 544 | 32.4% |

***Supplementary Table 3.*** Infection Prevalence Among Adult Ixodes Ticks Collected from Dogs by Treatment Group and Year

| **Year** | **Treatment Group** | **FlaB Positive** | **Total Ticks Tested** | **Proportion** |
|---|---|---|---|---|
| Baseline | TBRTV | _ | _ | _ |
| | Control | _ | _ | _ |
| 1 | TBRTV | 1 | 8 | 12.5% |
| | Control | 1 | 3 | 33.3% |
| 2 | TBRTV | 1 | 4 | 25.0% |
| | Control | 2 | 6 | 33.3% |
| 3 | TBRTV | 2 | 9 | 22.2% |
| | Control | 4 | 28 | 14.3% |
| 4 | TBRTV | 2 | 18 | 11.1% |
| | Control | 2 | 7 | 28.6% |
| Total | TBRTV | 6 | 39 | 15.4% |
| | Control | 9 | 44 | 20.5% |
